# Increasing network stability towards large food webs

**DOI:** 10.1101/2020.07.02.183723

**Authors:** Robert Veres, Zoltán László

## Abstract

Stability is a key attribute of complex food webs that has been for a long time in the focus of studies. It remained an intriguing question how large and complex food webs are persisting if smaller and simple ones tend to be more stable at least from a mathematic perspective. Presuming that with the increasing size of food webs their stability also grows, we analyzed the relationship between number of nodes in food webs and their stability based on 450 food webs ranging from a few to 200 nodes. Our results show that stability increases non-linearly with food web size based both on return times after disturbance and on robustness calculated from secondary extinction rates of higher trophic levels. As a methodologic novelty we accounted for food web generation time in the return time calculation process. Our results contribute to the explanation of large and complex food web persistence: in spite of the fact that with increasing species number the stability of food webs decreases at small node numbers, there is a constant stability increase over a large interval of increasing food web size. Therefore, in food web stability studies, we stress the use of food web generation times.

## Introduction

Importance of ecological networks has substantially increased in the last decades^1,2^. On one hand due to scientific advances in network theory^3^, on the other due to the increased need to understand the patterns behind persistence of living communities^4^. Because of significant decrease of biological diversity and threat of further acceleration of this decrease, the motivation of studying ecological network persistence and resilience has gained more importance than ever^5^.

The decrease of biological diversity is due to several reasons like human activities through habitat loss and fragmentation, and climatic change accelerated by large-scale human activities^6^. Climatic change will cause area shifts for numerous species, therefore whole communities will face extinctions and insertions^7^. These shifting species and communities will alter ecosystems, causing either shrinkage, or growth, whereas in some cases can lead to network collapse^8^.

Because collapses and extinctions of whole communities, stability characteristics as persistence and resilience of ecological networks became common study subjects^9^. Persistence describes the time at which communities will survive under changing environmental conditions^10^. Resilience describes the time which is needed by a system, in this case a community, to regain a stable structure similar to the one present before disturbances^10^. Stability of ecological networks raised intense and extended debates known as the complexity-stability debate^11–13^. The core question of the debate is: in which way may complex ecological networks persist, while reduced networks seems to be more stable^14,15^? The complexity and stability debate prevailed over the past five decades having presumably no simple resolution due to its complexity. Presumably several good resolutions may exist ^9,16,17^.

Complex ecological networks are predicted in theory to be more fragile than simple ones since complexity affects specie’s likelihood to be in stable equilibrium^18^. Moreover, variability of population densities over time also affects this community composition persistence^18^. But this complexity-fragility relationship seems to be not persistent in several cases, thus a general consensual agreement is lacking^9^. Spatiality may have a more pronounced stabilizing effect on complex food webs than it was predicted based on ecological theory^16,19^. Moreover, the complexity stability relationship seems to be lacking in empirical food webs since empirical food webs have non-random properties^17^.

However, before the 1970s, ecologists supported the idea that complex communities are more stable than simple ones, considering that natural communities develop into stable systems through successional dynamics^9^. Then, May’s (1972, 2001) work, which attracted also much criticism has shown that randomly assembled small communities are more stable that the large ones^21,22^. Since then it has been subsequently demonstrated that real world ecosystems differ significantly from randomly assembled model ecosystems taking into account their multifaceted structural patterns^17,22,23^

Data acquisition from real world ecosystems remained challenging also for the last decades, with a potential enhancement via molecular methods (e.g. metabarcoding or metagenomics)^24^. In recent studies adhering to the complexity-stability debate empirical ecosystems of different sizes are used, but the majority still handles mainly small (S<50, S = species number) food webs ^17^. Studies targeting large (S>100) food webs are relatively scarce^25^, therefore generalizations towards real world ecosystems are still questionable.

We analyzed resilience change of large food-webs through a wide size range of available empirical data to find support for early ecologist’s initial presumption that complex communities should be more stable than simple ones^9^ and to gain new insights to the realistic part of the complexity-stability debate. Our study hypothesis was that stability of food webs grows with complexity. To analyze this hypothesis, we chose network metrics based on different methodologies: i) the resilience measured by return times from the disturbance event^26^ and ii) the network robustness based on secondary extinction curves^27,28^. The chosen metrics lead to comparable results with both May’s approach and with more derived approaches, e.g. those used in many ecological recent studies. We formulated the following prediction: both food web resilience and robustness (stability) increases with network size (complexity) regardless of habitat type.

## Results

### Food-web descriptive statistics

Our analyzed data set comprised a total of 450 quantitative food webs with node numbers between 3 and 263. From the analyzed food webs a total of 85 were larger than 50 nodes (18.88%), while 34 were comprised more than 100 nodes (7.55%). Of the analyzed data sets 258 food webs were aquatic (57.33%), of which 206 were collected from freshwater, and 52 were from marine habitats. Of the remaining food webs 130 were terrestrial (28.88%) and 62 were mixed (13.77%). The mixed food webs comprised estuaries, swamps, shores and beaches; ecotone zones, boundaries of marine and terrestrial habitats.

All the used food webs were compared from a topological viewpoint using different network metrics to ensure that they are representative for the whole range of real food webs (Figure 1). The habitat types showed significant differences, except for the number of network nodes (S) (Gamma GLM: χ^2^=5.38, df=3, p=0.14). In the case of linkage density, food-webs of marine habitats had significantly higher linkage density than the others, while terrestrial had significantly smaller linkage density (Gamma GLM: χ^2^=43.02, df=3, p<0.001). For the smallest mean distance, the freshwater and terrestrial food-webs had smaller mean distances than the marine and mixed ones (Gamma GLM: χ^2^=22.47, df=3, p<0.001). In the case of connectance, food-webs of marine habitats had significantly higher connectance than the others, while terrestrial ones had significantly smaller linkage density (Gamma GLM: χ^2^=40.75, df=3, p<0.001).

**Figure 1.**
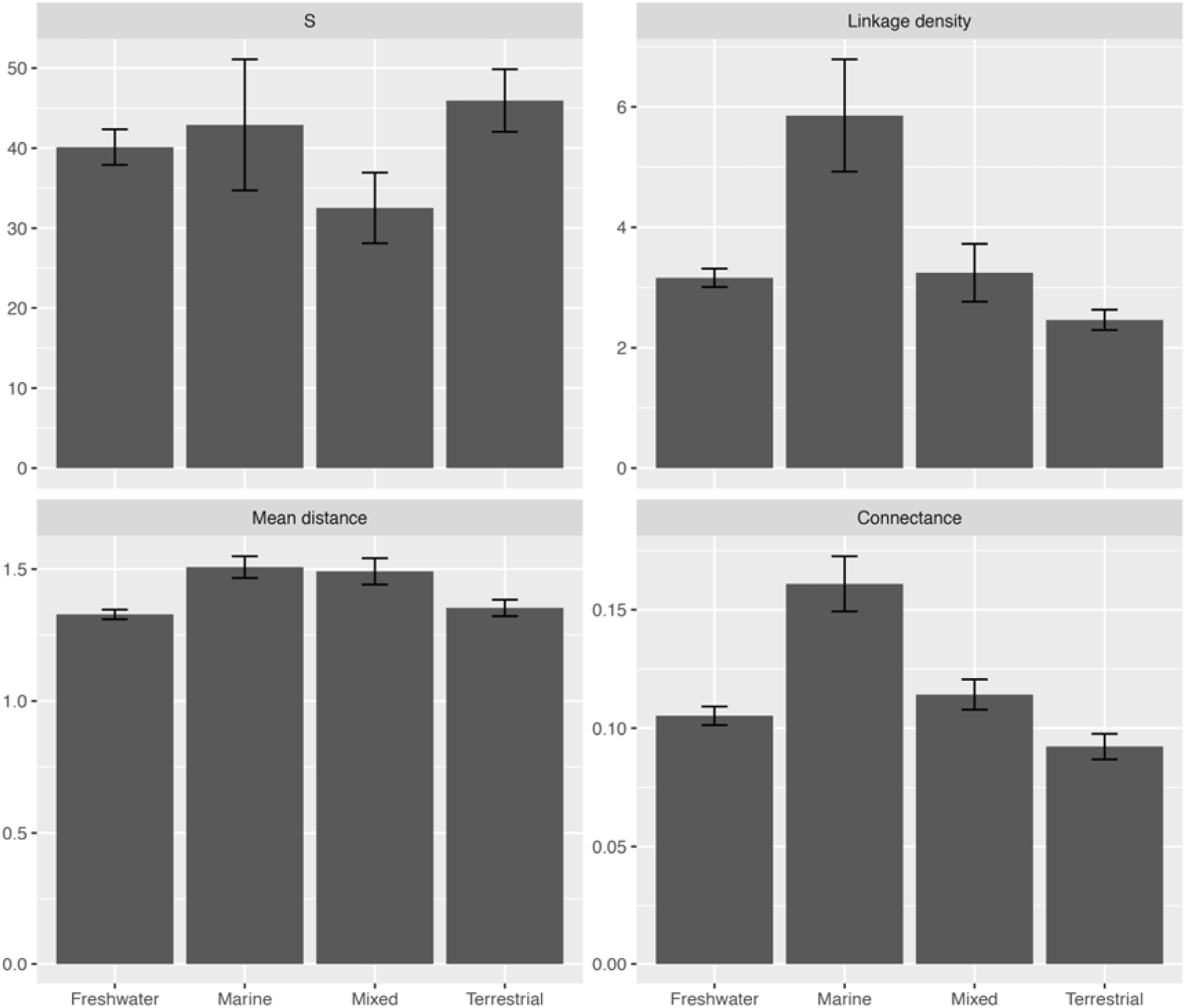
**Comparison of food-webs from different habitat types,** based on commonly used network metrics: node number (S), linkage density, smallest mean distance, connectance.

### Stability metrics

For the *return time* calculation, we performed Monte-Carlo simulations on the adjacency matrices of downloaded food-webs (Supplementary material S1 Table 1). For a single food-web, first we set the interaction strengths of nodes as defined by Pimm & Lawton^14^ Second, we randomly changed these interaction strengths 20,000 times, after which we calculated the leading eigenvalues of each randomized matrix. If a food web showed leading eigenvalues with negative real parts, then it was considered to converge to a local stability. If the leading eigenvalues had only positive real parts, then the food webs’ convergence to a local stability would not occur. We than took the return times as the negative reciprocal of leading eigenvalues with negative real parts. From these, we calculated the mode of their 5% trimmed, normalized frequency distribution, which was used as the representative return time of the respective food-web. We applied this procedure for every downloaded food-web. Analyzed food webs with positive leading eigenvalues (N=34) were excluded from the return time analysis.

To validate the use of normalized frequency distribution modes we also calculated the non-normalized frequency distribution modes of the 5% trimmed return times. The relationships between non-normalized and normalized return time modes with respect to food web size were opposite: non-normalized return time modes increased significantly, while normalized return times decreased significantly with food web size, regardless of ecotypes (Figure 2). However, the decline of normalized return times was most pronounced in marine ecosystems and weakest in the terrestrial ones (Table 1).

**Figure 2.**
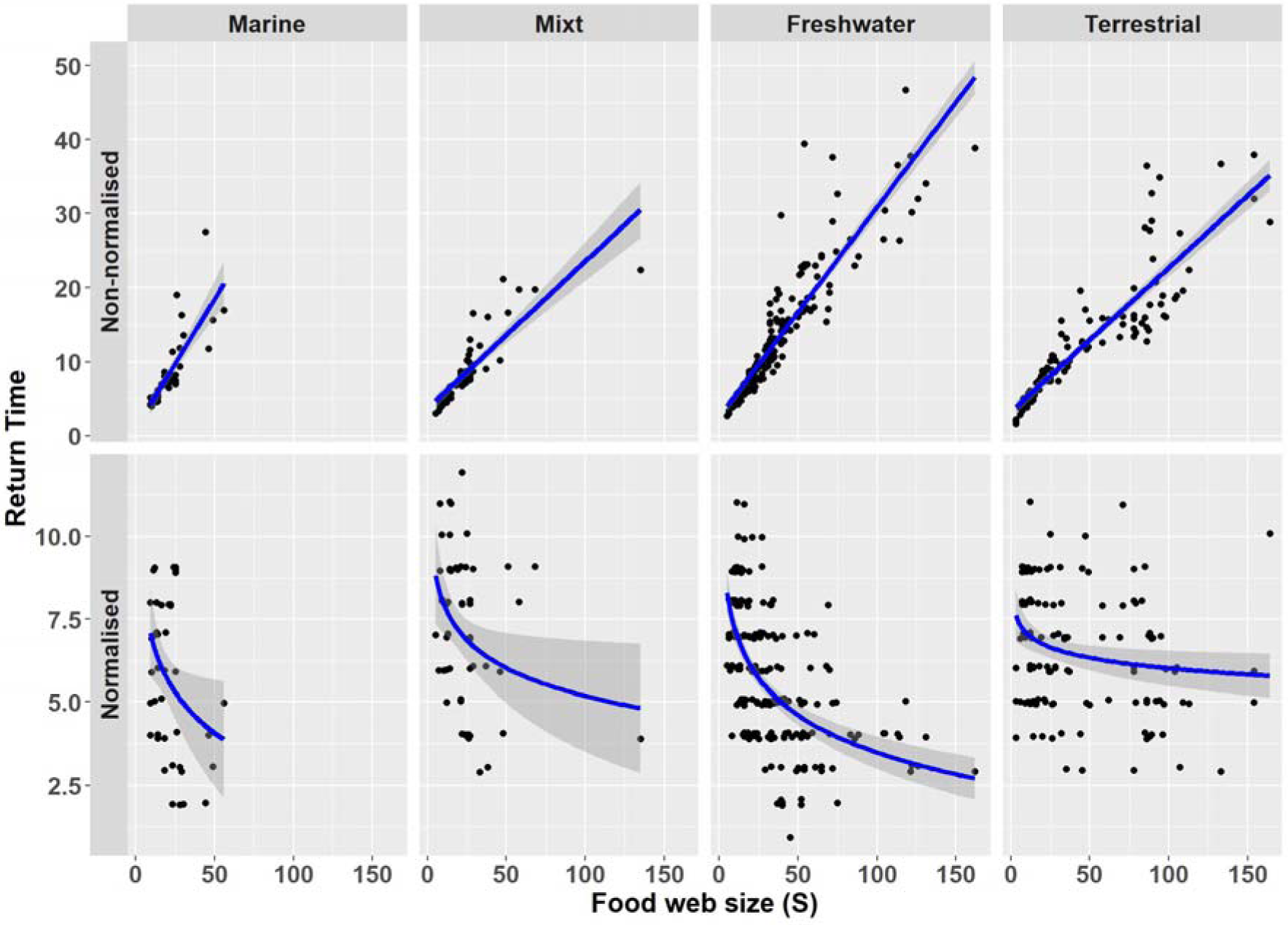
Relationships between non-normalized and normalized modes of return times (RT) and food web sizes. Non-normalized mean return times increased with food web size, while normalized return times decreased with food web size, regardless of ecotypes (N=416, marine: N=42, mixt: N=57, freshwater: N=190, terrestrial: N=127). RT coordinates are presented with a jitter effect.

**Table 1.** Opposite relationships between non-normalized and normalized mean return times with food web size.

|  |  | Non-normalized | Normalized |
| --- | --- | --- | --- |
| Marine | slope | 0.35 | -1.76 |
|  | SE | 0.04 | 0.73 |
|  | p | <0.001 | 0.02 |
| Mixt | slope | 0.20 | -1.21 |
|  | SE | 0.02 | 0.48 |
|  | p | <0.001 | 0.02 |
| Freshwater | slope | 0.28 | -1.61 |
|  | SE | 0.01 | 0.17 |
|  | p | <0.001 | <0.001 |
| Terrestrial | slope | 0.19 | -0.46 |
|  | SE | 0.01 | 0.16 |
|  | p | <0.001 | 0.01 |

For all analyzed networks regardless of ecotypes the relationship between non-normalized return times and food web sizes for the small size range (N=56) showed an increase on both linear and LOESS fits (Figure 3a). Also, for the whole food web size range (N=416) the non-normalized return time modes increased on both the linear and LOESS fits (Figure 3b). The linear relationship between non-normalized return time modes and small food web size (S range: 3-10 nodes, N=56 food webs) showed an increase in return time with the increasing size of food webs (linear LS: intercept=0.96, SE=0.22, t=4.45, p<0.001; slope=0.38, SE=0.03, t=14.27, p<0.001; F=203.68, p<0.01, N=54) (Figure 3a). For the whole food web size range (S range: 3-164, N=416) the linear relationship between non-normalized return time modes and food web size, also showed a continuous increase over the entire size range (linear LS: intercept=3.76, SE=0.34, t=11.08, p<0.001; slope=0.22, SE=0.01, t=32.19, p<0.001; F=1035.97, p<0.001, N=362) (Figure 3b).

**Figure 3.**
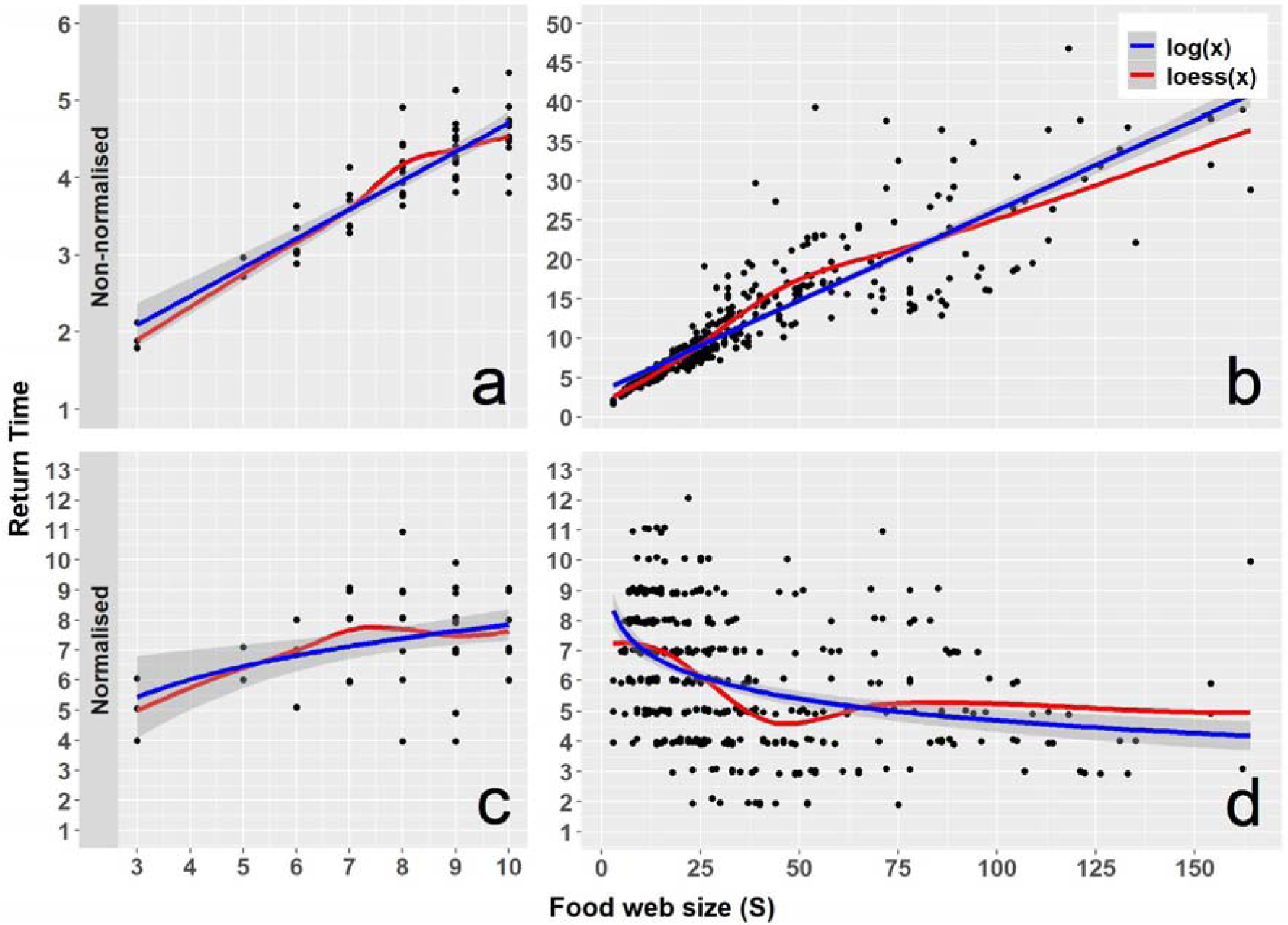
Relationship between return times and food web sizes. a) For small food web (N=56) size ranges, the non-normalized return time modes increased on both the log and LOESS fits. b) For the whole food web size range (N=416), the non-normalized return time modes increased on both the log and LOESS fits. c) For small food web (N=56) size ranges, the normalized return time modes increased on both the log and LOESS fits. d) For the whole food web size range (N=416), the normalized return time modes decreased continuously on the log fit and showed decrease, with a local increase in the LOESS fit. RT coordinates are presented with a jitter effect.

Again, for all analyzed networks regardless of ecotypes the relationship between normalized return times and food web sizes for the small size range (N=56) showed an increase on both log and LOESS fits (Figure 3c). While, for the whole size range (N=416), the normalized return time modes decreased continuously on both the log and LOESS fits (with exception for the aforementioned small food web size range of S=3-10) (Figure 3d). The logarithmic relationship between normalized return time modes and small food web size (S range: 3-10 nodes, N=56 food webs) showed an increase in return time with the increasing size of food webs (logarithmic NLS: intercept=3.23, SE=1.42, t=2.28, p=0.03; slope=2.00, SE=0.68, t=2.92, p=0.005; F=8.55, p=0.005, N=54) (Figure 3c). For the whole food web size range (S range: 3-164, N=416) the logarithmic relationship between normalized return time modes and food web size showed a continuous decrease over the entire size range (logarithmic NLS: intercept=9.78, SE=0.54, t=18.17, p<0.001; slope=-1.13, SE=0.15, t=-7.27, p<0.001; F=52.9, p<0.001, N=362) (Figure 3d).

Based on the LOESS weighted polynomial regression procedure for curve fitting, the normalized return time mode values begin to decrease drastically after the food web size of S=10, and after a slight increase around S∼50, it continued the decrease for the remaining size range (S_max_=164) (Figure 3d).

For the calculation of robustness, we used the area under the secondary extinction curve ^28^ also known as the attack tolerance curve ^27^. The tolerance curve shows how many extinctions (species loss) may occur at the pray level of a food web without reaching a total loss of the predator level. High values of robustness represent a more stable food web, where the drastic decrease in the number of predators occurs only after a high number of pray losses. For low robustness values, the drastic decrease occurs after just a few pray losses. Thus, a low robustness value represents an unstable food web. Robustness values were set between 0 and 1. Robustness values could not be calculated in case of two food webs, because of the small size and small number of connections. Thus, the robustness dataset contained N=448 analyzed food webs.

The logarithmic relationship between robustness and small food web size (S range: 3-10 nodes, N=52 food webs) showed an insignificant increase in robustness with the increasing food web size (logarithmic NLS: intercept=0.53, SE=0.09, t=5.84, p<0.01; slope=0.05, SE=0.04, t=1.26, p=0.22; ANOVA log(x): F=1.58, p=0.21, N=52) (Figure 5a). The logarithmic relationship between robustness and food web size showed a significant increase over the entire size range (S range: 3-164, N=448) (logarithmic NLS: intercept=0.49, SE=0.02, t=29.46, p<0.001; slope=0.07, SE=0.005, t=14.05, p<0.001; ANOVA log(x): F=197.28, p<0.001, N=448) (Figure 5b).

**Figure 5.**
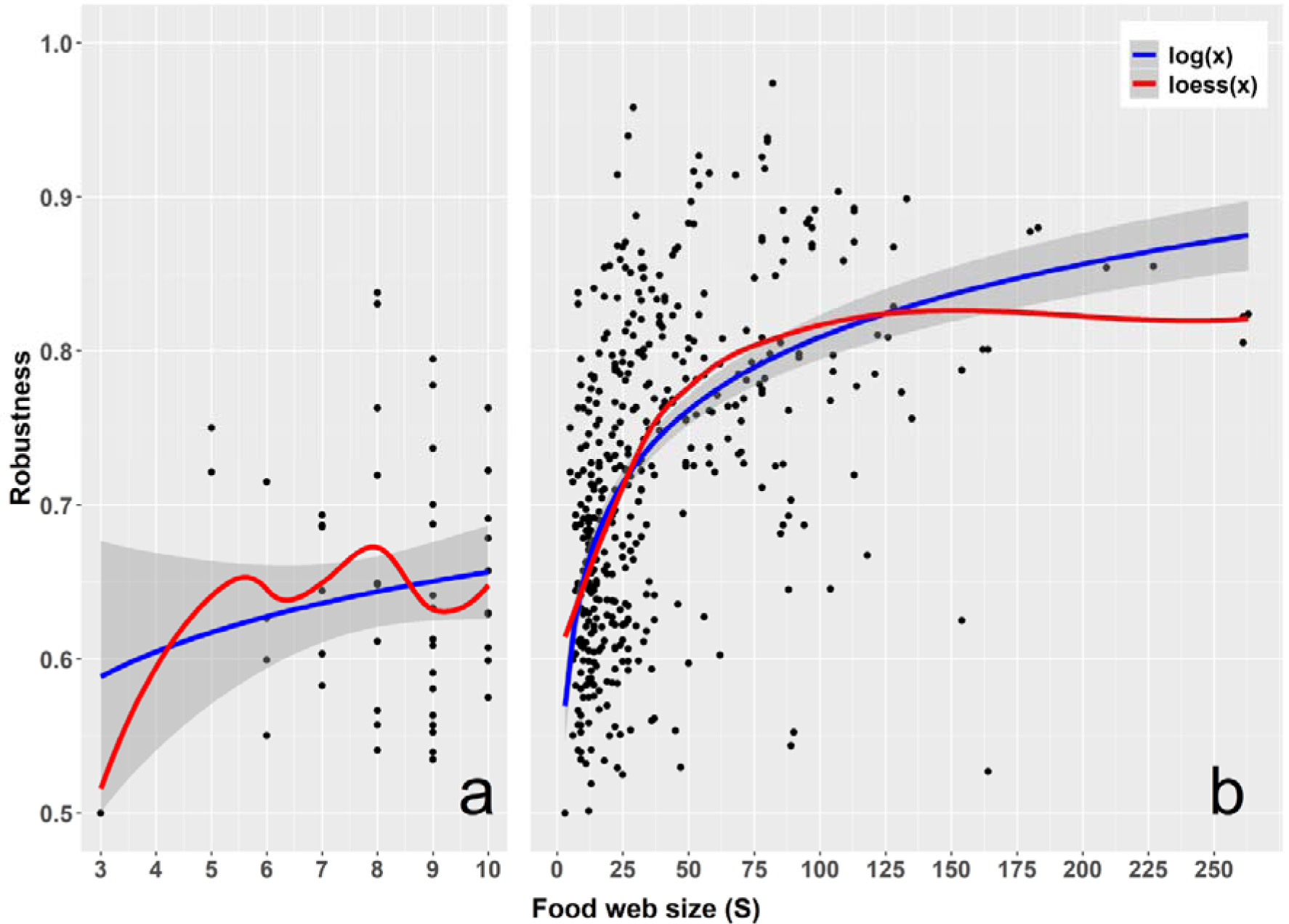
Relationship between robustness and food web sizes. a) For small food web (N=52) size ranges, the robustness increased on both the log and LOESS fits. b) For the whole food web size range (N=448), the robustness increased continuously on the log fit and showed also increase, with a local decrease in the LOESS fit.

Based on the LOESS weighted polynomial regression procedure for curve fitting the robustness values showed an increasing trend in the food web size of S=10 and, and showed again an increase for the remaining size range too (S_max_=263). The robustness values by habitat types also shows an increasing trend with the increasing food web size (Figure 6).

**Figure 6.**
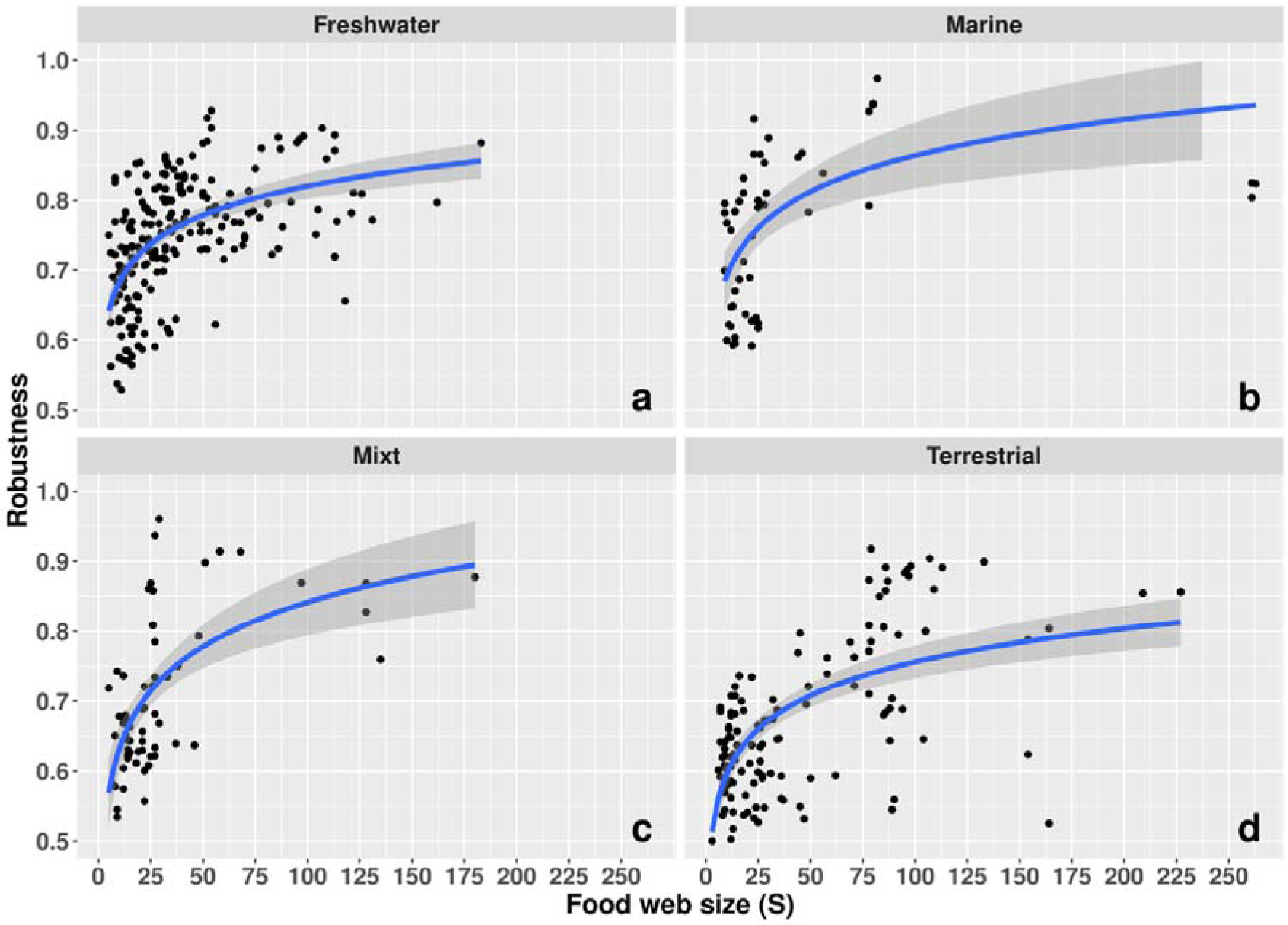
Relationship between robustness and food web sizes for habitat types. The robustness values increased with food web sizes in the four habitat types: freshwater (N=186), marine (N=47), mixed (N=56) and terrestrial (N=127).

## Discussion

Our study is one of the few analyses that have also included empirical systems with more than 200 species^9,16,17,19,29^. While we analyzed an extensive empirical data set, the majority of preceding studies on network stability have either investigated theoretical networks; or, if empirical data sets were used, these contained generally few and small networks^15,26,30,31^. To our best knowledge until now this is the largest analyzed empirical data set from the viewpoint of food web stability.

We analyzed the complexity-stability relationship in a large range of empirical food web sizes by return times and robustness. Our results led us to an assumption contrasting the presumption that stability should decrease with increasing food-web complexity: large food-webs have shorter return times than small ones if return-times are scaled (min-max normalized). The explanation for our assumption is based on the followings. Large food-webs have higher trophic levels which usually incorporate more large bodied species^32,33^ which have increased mean generation time^34,35^. Generation times affect food-web dynamics, having a significant impact on the return times^36^. Thus, when comparing different sized food-webs using their return times, a min-max normalized scaling is required for correction of the bias due to differences in generation time and their effects on return times. Such, we actually obtained via min-max normalization, generation time free return-times. Our results of generation-time free food-webs (Figure 3a) also support those in the literature: for networks with 10 or fewer nodes, the typical return time increases with node numbers^15,26^. This means that, in the small size range, larger systems take on average more time to return to a steady state after a random disturbance has occurred.

In terms of food web structure and other properties, the collected networks were diverse but similar enough to be comparable based on topological parameters. Based on the studied food webs, marine ecosystems differ from other habitats and averages over the entire dataset in several cases. These phenomena can be attributed to the characteristics of this habitat type. Our result is in concordance with other food web analyses results^35^. However, our results were consistent independently from metrics and habitat types. Regardless of habitat types, return times showed a decrease with increasing network size, and similar results were observed in terms of robustness. This has two implications: on one hand, the increase in stability towards large networks is non-habitat specific; on the other hand, an independent metric that works with other mathematical methods has produced the same result (robustness).

Using a straightforward metric such the secondary extinctions-based R, resulted an increase in stability with food-web size. By using normalized RT calculated from the 95% of their distribution, in contrast to May’s methodology where only the 0-150 interval RT were used, we found the same results as the R showed. Beyond this, we also found that May’s presumptions are valid both the normalized and 0-150 interval RT only for small food webs, while for larger ones a positive relationship between complexity and stability is the rule.

Robustness is used often for mutualistic and small food webs^1,27,28,37^. Our results confirms the results of Dunne et al.^38^: the robustness increases with the size of the network and with the resulting complexity. For estimating robustness, we applied their methods (estimation by secondary exclusions), and the size range of their networks (S =25-172) overlaps well with our data set, although the number of examined networks was relatively small (N=16).

Although we have confirmed that both robustness and return time are just as useful for describing network stability (the two approaches are mutually supportive), we recommend using return times as it can provide temporal information for stability. In addition, our results bring to light new, concise ideas about the “complexity-stability” relationship. Consistent with previous assumptions, the debate targeting the “complexity-stability” relationship will probably not be solved by a single, simple theory, but by a combination of several theories and approaches. Accordingly, the results we obtained could potentially bridge the gap between previous strong mathematical theories and conflicting empirical studies

The return time is an attractive metric from mathematical point of view. Stability studies based on the eigenvalues of system matrices have a long history both among mathematicians and physicists. As a result, computational techniques and theories based on it have become sophisticated and gaining popularity in other fields of research. This metric is also interesting because it can determine the amount of time a system needs to reach equilibria. It also provides an intuitive indication of stability. Since the initiating controversial results^15,20,26,39^, there has been an increasing number of studies based on return time metrics. Although the method is uniform, the results of studies are often contradictory. One of the significances of our results is that it demonstrated the stability of large networks in terms of return time and robustness, given the contradictions found in the literature. These have remained unresolved until now, due to the different analytical methods and used criteria^9,12^.

While we found a significant relationship between complexity and stability, Jacquet et al.^17^ showed that empirical food webs have several non-random properties that contribute to the lack of a complexity-stability relationship. However, in their study stability was dependent, not on network size, but on complexity. Moreover, while they analyzed 116 networks, their largest network size included only 50 species. It can be seen in Figure 2 that the increase in stability in our data appears around node number of 50, and the increase is constant for higher sizes based also on the LOESS curve. Other recent approaches^16,19^ attempted to explain the stability of large networks by spatial characteristics. Thus, our study is not comparable to these, but the methodology used are based on either return times or the use of Jakobian matrices which further may prove the effectiveness of these techniques. However, the negative correlation between stability and complexity was previously also contrasted with the constraint that longer food chains may have shorter return times than shorter ones if their energy flow is fast enough^40,41^.

Our approach in this way corrects the relationship between stability of food-webs and their size, by handling correctly the generation time effect on return times. This result brings a new insight to the understanding of the controversial relationship of the stability and complexity in food webs.

## Material and methods

### Data analysis

#### The data

Throughout the present study, we used a set of 450 empirical food webs compiled from a variety of online and publicly available sources. The majority of the food webs (∼80%) originated from www.globalwebdb.com, which is an online database housing a large set (359) of published empirical food webs. A part of the used food webs (51) are available through the ECOWeB database^42^. Another part of the food webs (26) are available from the ferencjordan.webnode.hu database. The compiled dataset contains a wide variety of food webs originating from different habitat types, spanning a wide size range (3-263 S), and having different levels of linkage density (0.67-36.02 links/species) (Supplementary material S1 Table 1).

We categorized each food web based on the habitat type it originated form, and defined 4 categories: aquatic habitats, which can be further broken down into freshwater and marine habitats; terrestrial habitats, which contain mostly non-aquatic species; and mixed habitat types, which are ecotone zones located on the boundary a terrestrial and different aquatic habitat types (ex. estuaries, marshes, swamps, shores and beaches, wetlands, bogs, mudflats, etc.).

We compared the compiled food webs from a topological viewpoint to ensure that the stability characteristics the food webs are representative, and are not a byproduct of the differences in topological structure. We calculated 4 widely used topological characteristics for each food web (Table 2, Supplementary material S1 Table 1), and used generalized linear models to assess the differences in the topology between the 4 different habitat types (Table 1).

**Table 2.** Topological metrics and their formulas used in the comparison of food webs.

| Name of the metric | Formula | Parameters |
| --- | --- | --- |
| number of nodes | $S$ | |
| linkage density | $\frac{L}{S}$ | $L$ – number of links |
| smallest mean distance | $\frac{1}{S \cdot (S - 1)} \cdot \sum_{i \neq j} d(v_i, v_j)$ | $d(v_i, v_j)$ - the shortest distance between $v_i$ and $v_j$ |
| connectance | $\frac{L}{S^2}$ | $L$ – number of links |

We studied the stability of food webs using two stability metrics: Return Time (RT) and Robustness (R). The main reason of choosing these metrics are the fundamental differences in the way they are calculated and the way they assess stability. By using metrics that are significantly different both in concept, and the way they are calculated, we reduce the risk of obtaining results biased by the used metrics.

#### Return Times (RT)

RT are derived from the linear stability analysis of a food web’s interaction matrix^43^. We derive such a matrix by the linearization of an *N*-species system of Lotka–Volterra equations at an equilibrium point. Each entry *a*_*ij*_ of the interaction matrix quantifies the change in population growth rate of species *i* caused by a small perturbation in the abundance of species *j* around equilibrium abundances. Such an interaction matrix can be viewed as the Jacobian matrix of the system, and it describes each species abundance dynamic over continuous time. We can determine the (local) stability of the equilibrium of the system by looking at its eigenvalues derived from the Jacobian matrix. If the leading eigenvalue (the eigenvalue with the largest real part) of the system has negative real parts, the equilibrium is stable (small perturbations at the equilibrium dampen over time). If the leading eigenvalue of the system has positive real parts, the equilibrium is unstable (small perturbations at the equilibrium are amplified over time). If an equilibrium is stable, we can use the negative inverse of the real part of the leading eigenvalue to determine the time needed for the system to return to a state of equilibrium after a small perturbation. The aforementioned time required to return to the equilibrium is the RT^9,15,20,39^. The aim of our study was not to demonstrate the validity of May’s stability metric^15,20^ and our analyses does not involve this stability criterium since its validity is debated^44^.

Throughout the simulations, we consider each food web to be at a state of local equilibrium. Default interaction strengths for each Jacobian matrix were set based on Pimm & Lawton^26^. Values in the upper triangle of the matrix (predator effect on prey) were set to −1 and values in the lower triangle of the matrix (prey effect on predator) were set to +0.1. The diagonals of each matrix (intraspecific interactions) were also set to −1. We simulated perturbances by multiplying each non zero element of the Jacobian matrices by random numbers drawn from a U(0, 1) distribution. After the randomization we calculated the leading eigenvalues, and determined their signs. We calculated the RT for each case in which leading eigenvalues with negative real pats were produced. We repeated these steps for each food web 20,000 times. After removing the food webs that did not produced leading eigenvalues with negative real parts, we obtained with a set of 416 food webs.

After performing the simulations, we get a number of RT between 0 and 20,000 for each food web, with values ranging from ∼1 to >2e+09 (largest RT encountered). This large range raises a potential problem. Values from the higher end of the RT range could be too big to be applicable in real life (it would take too long for the system to return to a state of local equilibrium to be applicable). Pimm & Lawton^26^ deal with this large range by excluding all RT that are greater than 150, and working only with values that are smaller than this threshold. This method is valid for small food webs (chains), but leads to another problem: while all food webs produce right skewed distributions of RT, and some of the modes (and thus the majority of the values) are very close, or even past this threshold, removing values close to the original (non-trimmed and non-normalized) modes of the distributions, leads to an artificial shift toward higher values. We tackle this problem, by performing a 5% trim at the higher end of our RT range. The 5% trim is enough to get rid of the long tail of the distribution with the extreme values, while keeping the most frequent RT values around the modes intact.

Another correction we used for each food web is connected to their generation time. If two food webs have the same size, but differ in mean generation time (ex. two food webs with 10 nodes, one comprised by herbs, insects and insectivorous birds while the other by trees, mammal herbivores and predators), their RT after perturbations will be proportional to their mean generation time. This means, that food webs with larger mean generation time will have on average lager RT values. To assess the relationship between RT and food web size, we had to correct for mean generation time. To correct for the mean generation time of the food webs, we performed min-max normalizations on the RT of each food web. By doing so, we scaled the RT range of each food web to [0, 1], making the distributions directly comparable.

We determined the representative RT of a food web, by creating bins on the [0, 1] interval in increments of 0.01 (a total of 100 bins in the case of each food web), and counting the number of RT within each bin. We took the mode of the RT distributions for these bins (a 1% interval) as the representative RT (the 1% of the most frequent RT in the case of a food web). Only in 7 cases, we obtained two modes for the RT distributions. In these cases, we chose the modes with the higher RT values.

Further sensitivity analyses targeting the construction of random Jacobian matrices could lead to a better understanding of stability dynamics, by drawing interaction strength multipliers from other distributions that U(0, 1), and by allowing correlated interactions strengths.

#### Robustness (R)

The main reason for using R was to have an independent stability measure that could be used to compare and test the results from the RT analysis. R is appealing, because is a simple stability metric, which can be easier interpreted than RT.

Interactions within a food web can be treated as bipartite interactions, where we consider the interactions of pairs of species (a predator and a prey) separately. This approach can be useful because by treating these systems as bipartite, we can utilize metrics developed to characterize bipartite system stability. One such metric is the secondary extinction rate (SER) of the upper level. In this case, predators are in the upper, while prey are in the lower level. For longer food chains, intermediate species are included separately as both predator and prey. To determine the SER of the upper levels, we randomly remove species one by one from the lower levels, and count the number of predators that remain without prey at the end of each step. It is hypothesized that such predators would become extinct due to lack of prey, and are excluded from the simulation from that point. This process is repeated until no species remain in the system^27,28^.

If we plot the number of surviving species after the extinction steps, we get a so-called attack tolerance curve that characterizes the survival of the system^27^. This approach can provide a good insight into the potential extinction cascades that can be triggered by the accidental extinction of certain species within a given network. SER can be modeled by applying spline interpolation to the attack tolerance curve using the function f(x) = 1-x^*a*^, where the exponent *a* represents the secondary extinction rates^28^.

The R values are derived from the SER^27,28^, and are defined as the area under the attack tolerance curve. We performed min-max normalizations on the x and y axes so that the values fall between 0 and 1, and thus the area under the attack tolerance curve (R) also falls within this range. This is useful because intuitively, an R value around 0 characterizes an unstable, less robust system, while a value around 1 characterizes a stable, much more robust system. This is because for low R values to occur, the number of prey species that are needed to be removed for mass extinctions to occur is low. This means, that the system will fall apart after a relatively low number of extinctions, indicating that the system is fragile. The opposite is true in the case of high R values: a larger number of extinctions are needed to occur for the system to fall apart, thus indicating that the system is more stable^27,28^.

SER simulations were performed 1000 times for each individual food web, and the average R value was calculated and used for the analysis in the case of each food web.

### Software and hardware

Statistical analysis and simulations were performed using the R statistical environment^45^. The GLM’s were evaluated using the “car” package^46^. Robustness simulations were done using a modified (parallelized) version of the “*robustness*” function from the “bipartite” package^47^. SQLite databases were used for storing and manipulating large data tables produced by simulations. Simulations were conducted on “Kotys” server of the BabelJ Bolyai University^48^.

## Acknowledgements

We are thankful to Joel E. Cohen for its valuable and critical comments on the manuscript. We thank to the “Kotys” cluster management team at the BBU Cluj for technical support during simulations. We also thank the valuable comments of several colleagues on our analyses and manuscript draft.

